# Novel determinant of antibiotic resistance: a clinically selected *Staphylococcus aureus clpP* mutant survives daptomycin treatment by reducing binding of the antibiotic and adapting a rod-shaped morphology

**DOI:** 10.1101/2023.03.06.531458

**Authors:** Lijuan Xu, Camilla Henriksen, Viktor Mebus, Romain Guérillot, Andreas Petersen, Nicolas Jacques, Jhih-Hang Jiang, Rico J. E. Derks, Elena Sánchez-López, Martin Giera, Kirsten Leeten, Timothy P. Stinear, Cécile Oury, Benjamin P. Howden, Anton Y. Peleg, Dorte Frees

## Abstract

Daptomycin is a last-resort antibiotic used for treatment of infections caused by Gram-positive antibiotic-resistant bacteria such as methicillin-resistant *Staphylococcus aureus* (MRSA). Treatment failure is commonly linked to accumulation of point mutations, however, the contribution of single mutations to resistance and the mechanisms underlying resistance remain incompletely understood. Here we show that a single nucleotide polymorphism (SNP) selected during daptomycin therapy inactivates the highly conserved ClpP protease and is causing reduced susceptibility of MRSA to daptomycin, vancomycin, and β-lactam antibiotics as well as decreased expression of virulence factors. Super-resolution microscopy demonstrated that the improved survival of the *clpP* mutant strain during daptomycin treatment was associated with reduced binding of daptomycin to the septal site and diminished membrane damage. In both the parental strain and the *clpP* strain, daptomycin inhibited the inward progression of septum synthesis eventually leading to lysis and death of the parental strain while surviving *clpP* cells were able to continue synthesis of the peripheral cell wall in the presence of 10 × MIC daptomycin resulting in a rod-shaped morphology. To our knowledge, this is the first demonstration that synthesis of the outer cell wall continues in the presence of daptomycin. Collectively, our data provide novel insight into the mechanisms behind bacterial killing and resistance to this important antibiotic. Also, the study emphasizes that treatment with last-line antibiotics is selective for mutations that, like the SNP in *clpP*, favor antibiotic resistance over virulence gene expression.

**IMPORTANCE:** The bacterium *Staphylococcus aureus* is a leading cause of life-threatening infections and treatment is challenged by the worldwide dissemination of methicillin-resistant *Staphylococcus aureus* (MRSA) that are multi-drug resistant. Daptomycin, a cell membrane-targeting cationic lipopeptide, is one of the few antibiotics with activity against MRSA, however, the killing mechanism of daptomycin and the mechanisms leading to resistance are not fully understood. Here we show than an MRSA strain, isolated from the blood of a patient treated with daptomycin, has acquired a mutation that inactivates the ClpXP protease resulting in increased resistance to several antibiotics and diminished expression of virulence genes. Super resolution microscopy showed that the mutant avoids daptomycin-elicited killing by preventing the binding of the antibiotic to the septal site and by growing into a rod-shaped morphology. In summary, this study discloses new perspectives on the mechanism of killing and the mechanism of resistance to an antibiotic of last resort.

## Introduction

*Staphylococcus aureus* is an opportunistic pathogen responsible for localized skin infections and severe illnesses, such as bacteremia, sepsis, osteomyelitis, and infective endocarditis (1). Treatment of *S. aureus* infections is challenged by the dissemination of methicillin-resistant *S. aureus* (MRSA) that in 2019 were associated with more than 100,000 deaths and 3.5 million disability-adjusted life-years worldwide (2, 3). The recommended treatment for invasive MRSA infections includes the last-resort antibiotics, daptomycin and vancomycin (4), however, *S. aureus* strains with decreased susceptibility to daptomycin and cross-resistance to vancomycin, and vice versa, have emerged during treatment (5–10). The development of resistance during therapy is a serious threat as it severely compromises treatment options, however, the molecular mechanisms underlying decreased susceptibility to daptomycin remain incompletely understood.

Daptomycin is a calcium-dependent lipopeptide antibiotic with rapid bactericidal activity against a broad range of Gram-positive bacteria. The mode of action of daptomycin has been a matter of debate as excellently summarized in a recent review (11). In a prevailing model, daptomycin in complex with Ca^2+^ functions like a cationic antimicrobial peptide that after binding to negatively charged phosphatidylglycerol kills Gram-positive bacteria by perturbing the integrity of the cytoplasmic membrane (11–13). However, daptomycin was also proposed to directly target the cell wall biosynthesis machinery (14–16). These apparently inconsistent results were reconciled by a recent study showing that daptomycin specifically targets undecaprenyl-coupled cell wall precursors at the division septum and that this initial binding is followed by massive membrane rearrangements and delocalization of peptidoglycan synthesis (17).

Daptomycin resistance (typically referred to as non-susceptibility) is often associated with genetic changes in the biosynthesis of membrane lipids that decrease daptomycin binding to, or penetration of, the cytoplasmic membrane (18–20). Additionally, mutations in genes encoding two-component systems controlling processes related to cell wall stress (*walKR*, *vraRS,* or *graRS*), quorum sensing (*agr*), and in *rpoB* and *rpoC* encoding subunits of the RNA polymerase, have consistently been reported in isolates collected after treatment failure (11; 21-23). Interestingly, vancomycin treatment has selected for mutations in the same set of genes, emphasizing that reduced susceptibility to vancomycin and daptomycin may be achieved by overlapping mechanisms (5-7; 24-26). The affected genes encode pleiotropic regulators controlling expression of large sets of genes and it has proved challenging to determine the downstream effectors improving tolerance to antibiotics.

The ClpXP protease is a highly conserved serine protease that is critical for protein turnover in bacteria and mitochondria (27). The ATP-dependent unfoldase ClpX recognizes substrate proteins and feeds them into the proteolytic chamber formed by 14 ClpP protease subunits (28). In *S. aureus*, ClpXP is essential for virulence in both systemic and abscess animal models of infection (29–32). Nonetheless, mutations in the *clpP* and *clpX* genes have on multiple occasions been identified in *S. aureus* strains isolated from patients undergoing treatment with daptomycin or vancomycin, raising the question of how *S. aureus* benefit from mutating the ClpXP protease *in vivo* (8,26,33–35). To answer this question, we here characterized a clinical MRSA isolate collected after daptomycin treatment failure that harbored mutations in c*lpP* and *rpoB* and examined how each of these mutations contribute to phenotypic changes. We show that the mutation in *clpP* eliminates ClpP activity and confers well-described *clpP* phenotypes such as diminished expression of virulence factors and augmented resistance to β-lactam antibiotics. Further, we for the first time demonstrate causality between inactivation of ClpP and reduced susceptibility to daptomycin and vancomycin and show that inactivation of the ClpXP protease improved survival in the presence of high, therapeutic concentrations of daptomycin. Super-resolution microscopy revealed that whilst daptomycin synchronized wild-type *S. aureus* cells in a stage of early septum synthesis that over time lose viability, the better survival of *clpP* cells was associated with decreased daptomycin binding, improved membrane integrity, and a shift in cell wall synthesis from the septal site to the peripheral wall resulting in cells adopting a rod-shaped morphology.

## Results

### Construction of *S. aureus* strains harboring single mutations in *clpP* (G_281_A) or *rpoB* (C_1430_A)

Two isogenic sequence type 5 MRSA strains isolated from the bloodstream of a patient with bacteremia before and after daptomycin treatment failure, respectively, A9781 and A9788 (for the ease of presentation denoted “SADR-1” and “SADR-2”) were previously characterized (8, 36). The pleiotropic phenotypic changes in SADR-2, as compared to SADR-1, were originally attributed to a SNP in the *rpoB* gene resulting in an A477D substitution in RpoB. However, re-sequencing of the SADR-2 genome revealed an additional SNP in the *clpP* gene (37). To elucidate the contribution of each of the two identified SNPs, *clpP* (G281A) and *rpoB* (C1430A) to the SADR-2 phenotypes, the two mutations were separated as described in Table 1. In short, SADR-1 derivatives harboring only the SNP in *rpoB* were constructed by either introducing the mutant *rpoB* allele into SADR-1 by allelic replacement (SADR-1*^rpoB_^*^1^ and SADR-1*^rpoB_^*^2^), or, by restoring wild-type *clpP* in SADR-2 resulting in SADR-1*^rpoB_^*^3^ and SADR-1*^rpoB_^*^4^ (27). With these four strains in hand, we could now unambiguously determine which of the SADR-2 phenotypes that are associated with expression of the RpoB_A477D_ variant. Moreover, the phenotypic changes not linked to the mutant *rpoB* allele could then be assigned to the SNP in *clpP*. To strengthen our conclusions, the G281A allele of the *clpP* gene was additionally introduced into the chromosome of SADR-1 by allelic replacement (37). We were, however, unable to obtain strains that deviated from SADR-1 only by the SNP in *clpP*, and therefore two derivatives (SADR-1*^clpP^*^_1^ and SADR-1*^clpP^*^_2^) harboring additional SNPs were used to back up our conclusions (Table 1).

**Table 1.**
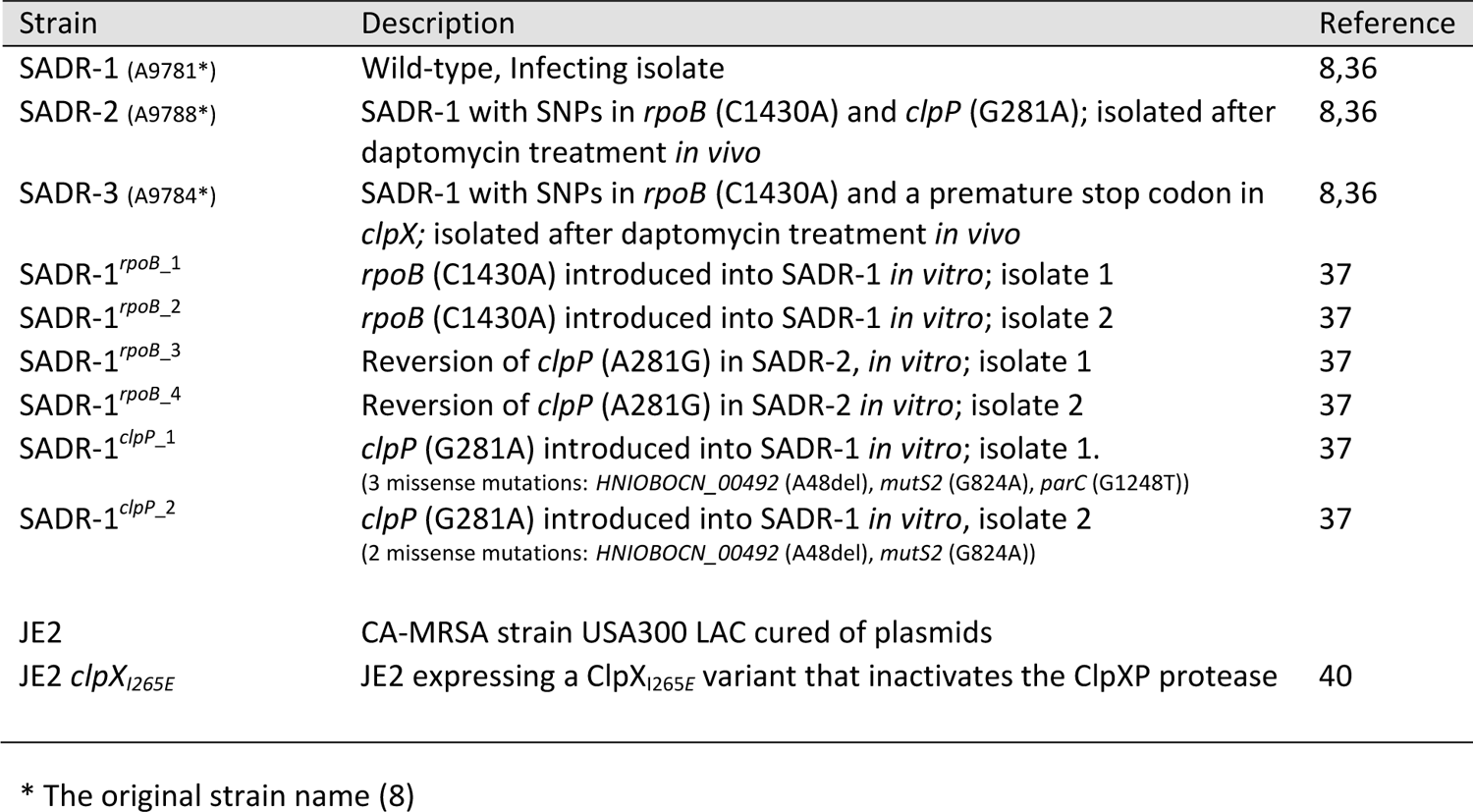
Strains used in this study

### The ClpP_G94D_ variant confers heat sensitivity, diminished cell size, reduced expression of virulence genes, and Sle1 accumulation

SADR-2 displayed a number of phenotypes that are typical for *S. aureus* mutants lacking ClpP activity, including heat sensitivity, diminished cell size, and decreased expression of virulence factors such as protein A, hemolysins and extracellular proteases (29, 36, 38–41). These phenotypes were originally attributed to the *rpoB* mutation but repeating these assays with the SADR-1*^rpoB^* and SADR-1*^clpP^* single mutants, clearly showed that these phenotypes are linked to the SNP in *clpP*, indicating that the ClpP_G94D_ variant is non-functional (Fig. 1 A-C). To strengthen this conclusion, we monitored the cellular levels of the Sle1 cell wall amidase that is degraded by ClpP (42, 43). ClpP does not interact with substrates directly, and in order to degrade proteins it needs to associate with one of two Clp ATPases that are responsible for substrate recognition and unfolding (40). In the case of Sle1 degradation the Clp ATPase, ClpX, is responsible for substrate recognition (42, 43), and in this experiment we included the isogenic SADR-3 isolate that cannot form the ClpXP protease due to a frameshift mutation in *clpX* abolishing ClpX synthesis (36). As seen in Fig. 1D, Sle1 accumulated to the same extent in SADR-3 and SADR-1 cells expressing the ClpP_G94D_ variant, supporting that theG94D substitution eliminates proteolytic activity of ClpP. In support here off, the substituted glycine residue, G94, localizes near the active site serine (S98) and is highly conserved in ClpP subunits from different kingdoms (human, plants, and bacteria) – Fig. 1E. We conclude that SADR-2 has acquired a mutation in *clpP* that abolishes ClpP activity resulting in heat sensitivity, diminished cell size, and reduced expression of virulence factors in SADR-2.

**Fig. 1.**
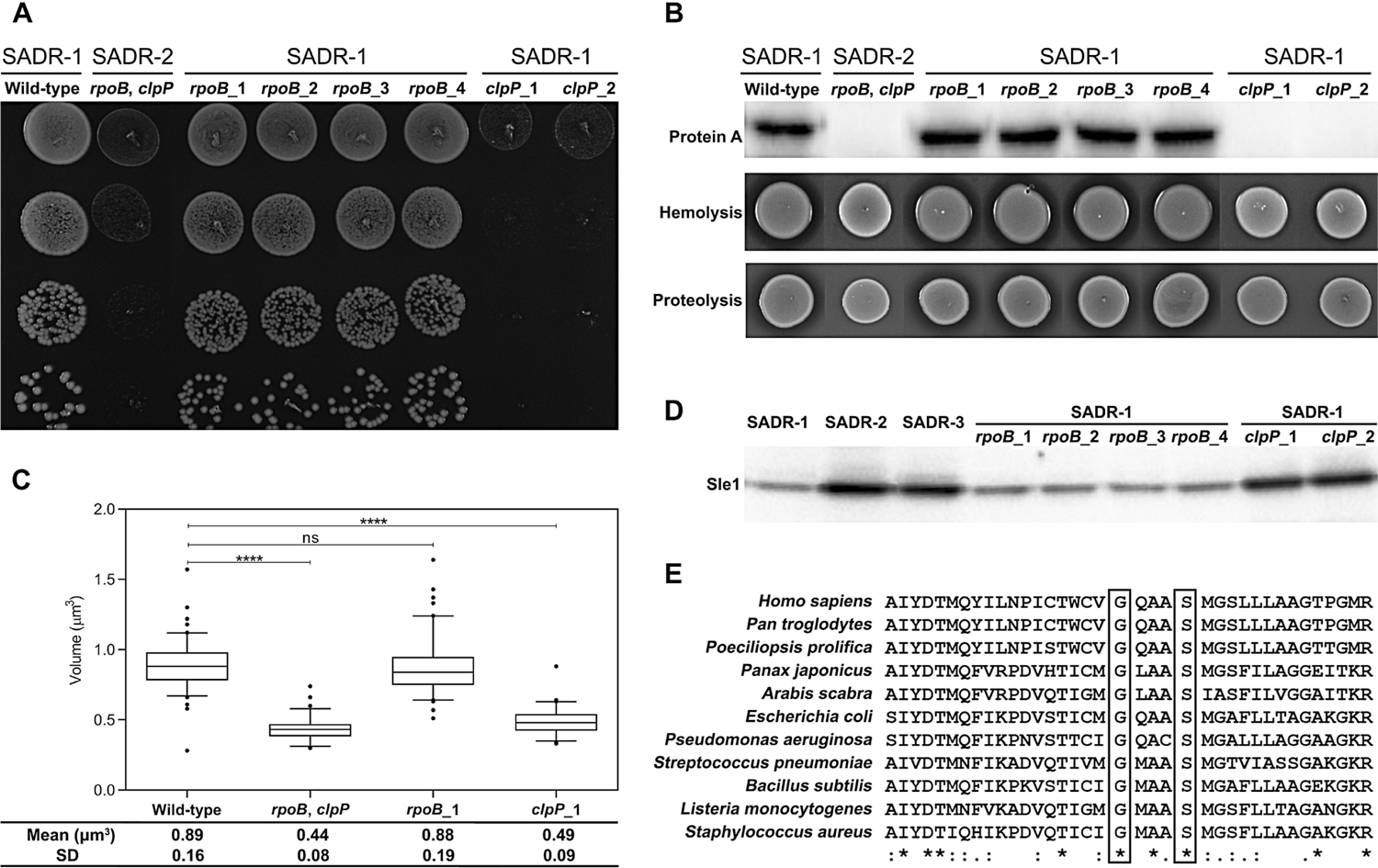
The G94D substitution in ClpP confers heat sensitivity, decreased expression of virulence, and diminished cell size. (A) Heat sensitivity was examined by a spot titer assay: aliquots (10 μl) of serial 10-fold dilutions of exponential cells grown at 37°C were spotted on TSA and incubated for 24 h at 42°C. (B) Cellular levels of protein A in late exponential growth phase were determined by Western blotting (top panel). Hemolytic activity (middle panel) and extracellular proteolytic activity (bottom panel) were detected as clearing zones on plates containing 5% calf blood or 10% skimmed milk, respectively. (C) The average cell volume of indicated strains was estimated in cells grown exponentially at 37°C (****, p < 0.0001 versus SADR-1). (D) The Sle1 levels were determined in whole cell extracts derived from cultures of the indicated strains in late exponential growth phase by western blotting. (E) ClpP sequences from different organisms were aligned using MUSCLE (MUSCLE < Multiple Sequence Alignment < EMBL-EBI). Fully conserved (identical) amino acids residues are marked by asterisks, similar residues are marked with colons (:). The substituted glycine (G94) and the active site serine (S98) are shown in boxes.

### The ClpP_G94D_ variant decreases susceptibility to daptomycin, vancomycin, and oxacillin

Compared to SADR-1, SADR-2 has decreased susceptibility to daptomycin, vancomycin, oxacillin, and rifampin – antibiotics belonging to four different classes of antibiotics (36). To examine if the decreased antibiotic sensitivity is caused by the *clpP* or the *rpoB* mutations, we determined the MICs of these antibiotics for the single- and double mutants derived from SADR-1 (Table 2).

**Table 2.**
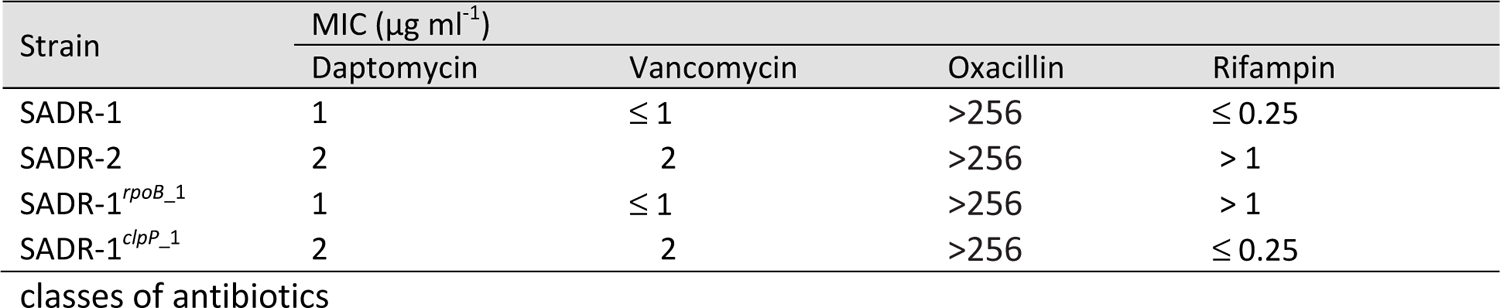
Antibiotic susceptibility of SADR-1 and the corresponding derivatives to four different

Consistent with previous results, the daptomycin MIC increased two-fold when going from SADR-1 to SADR-2 (36). A similar increase in daptomycin MIC was conferred by introduction of the *clpP* SNP into SADR-1, while introduction of the mutant *rpoB* allele did not change the daptomycin MIC (Table 2). Similarly, the sole introduction of the mutated *clpP* allele in SADR-1 increased the vancomycin MIC two-fold, while the vancomycin MIC was unchanged by expression of the RpoB_A477D_ variant. Finally, the single point mutation in *rpoB* increased the rifampin MIC to the level measured for SADR-2, while the mutation in *clpP* did not impact rifampin MIC (Table 2). This finding is consistent with reports showing that an A477D substitution in RpoB causes resistance to rifampin (44, 45).

The presence of subpopulations with higher resistance than the main population, the phenomenon of hetero-resistance, has come into focus as a cause of treatment failure (46). Hence, antibiotic susceptibility was additionally determined by performing population analysis profiles (PAPs). The oxacillin MIC of SADR-1 was already very high (>256 µg ml^-1^, Table 2), however, in PAP analyses, SADR-1 displayed heterogeneous resistance to oxacillin with only a small fraction of SADR-1 cells being capable of forming colonies at oxacillin concentrations exceeding 8-16 µg ml^-1^ (Fig. 2). Consistent with published data, SADR-2 cells are homogeneously resistant to high concentrations of oxacillin (36), and a similar phenotype was observed for SADR-1*^clpP^*^_1^, demonstrating that inactivation of ClpP converts SADR-1 from a heterogeneously resistant to a homogeneously, highly resistant MRSA strain. However, the SNP in *rpoB* also increased resistance to oxacillin at the population level as evidenced by the right-shift of the curve (Fig. 2). In accordance with the higher vancomycin MIC of SADR-2, a right-shift of the curve was observed in the PAP analysis. Importantly, a similar right-shift was observed for SADR-1 expressing the ClpP_G94D_ variant demonstrating that inactivation of the ClpP protease enables a larger fraction of cells to form colonies at the higher vancomycin concentrations (Fig. 2). The PAP analyses, therefore, support that the decreased sensitivity to vancomycin is associated with the SNP in *clpP*. The PAPs in Fig. 2, however, also revealed a small right shift for the strains only carrying the mutant *rpoB* allele supporting a previous study showing that an A477D substitution in RpoB contributes to the decreased susceptibility to daptomycin and vancomycin at the population level (45).

**Fig. 2.**
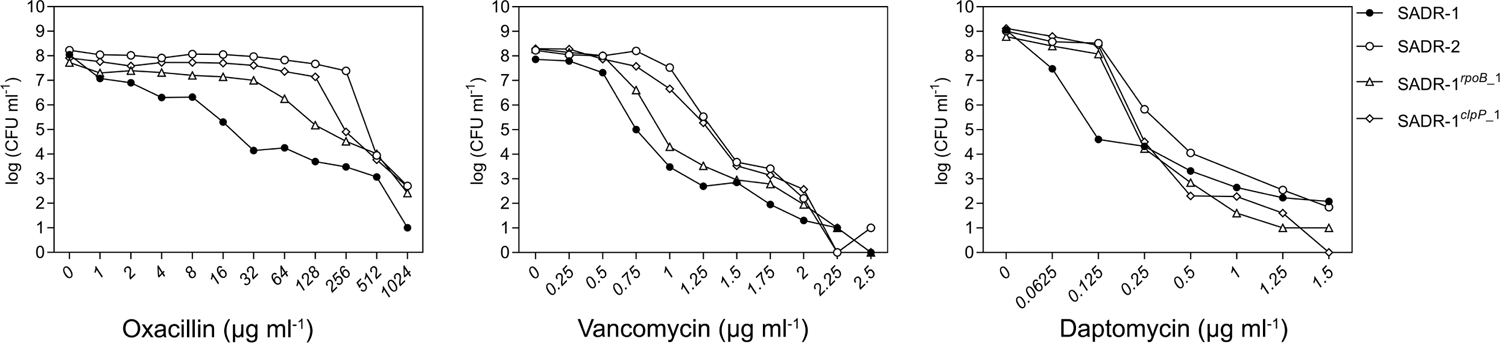
Population analysis profiles show that inactivation of *sle1* renders *S. aureus* cells homogenously susceptible to vancomycin and daptomycin. CFU ml^-1^ was determined after plating on increasing concentrations of vancomycin, oxacillin, or daptomycin (+ 50 µg ml^-1^ CaCl2) as indicated. Representative data from three individual experiments are shown.

In conclusion, inactivation of ClpP accounts for the increase in daptomycin, and vancomycin MICs, and increased subpopulations with higher resistance to vancomycin and oxacillin, however, the *rpoB* allele causes a minor increase in tolerance to daptomycin, vancomycin, and oxacillin at the population level.

### Inactivation of ClpXP promotes *S. aureus* survival at high daptomycin concentrations

SADR-2 was selected during daptomycin therapy *in vivo* and we next asked if inactivation of ClpP also promotes survival of *S. aureus* cells when exposed to high concentrations of daptomycin (10 × MIC) that mimic the clinically relevant dose. In this experiment, we like others used a high inoculum of bacteria (2 × 10^8^ CFU ml^−1^), as daptomycin is licensed for treatment of infective endocarditis which is associated with high bacterial loads (47, 48). In this assay, most SADR-1 cells were killed rapidly within 2 h (3-log reduction in CFU counts) in the presence of 20 µg ml^-1^ daptomycin while SADR-2 cells survived markedly better (killing reduced ∼100-fold), Fig. 3A. Interestingly, the SNP in *clpP* alone promoted survival, while introduction of the *rpoB* mutation alone had no effect (Fig. 3A). To determine if ClpP proteolytic activity impacts *S. aureus* daptomycin survival across different clonal complexes we included the JE2 model strain that belongs to the clinically important MRSA USA300 clone (CC8). Interestingly, daptomycin survival was improved 100-fold in JE2 cells expressing a ClpX_I265E_ variant that cannot associate with the ClpP proteolytic subunits to form the ClpXP protease, Fig. 3B (40). Together, our results show that inactivation of the ClpXP protease is associated with higher daptomycin survival across different clonal complexes.

**Fig. 3.**
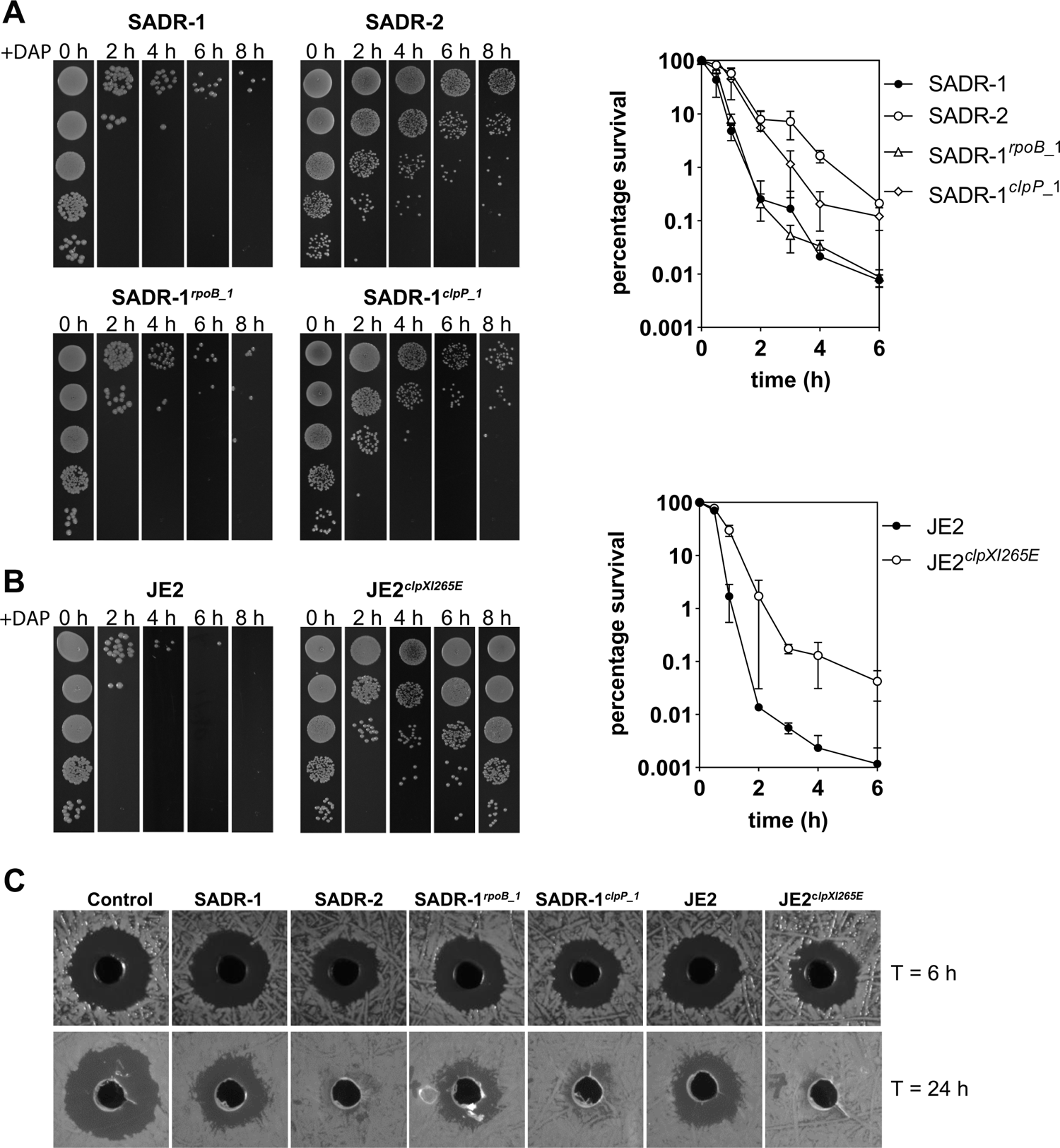
Inactivation of ClpP promotes survival during daptomycin treatment. Stationary cells (∼2 × 10^8^ CFU ml^-1^) were resuspended in TSB supplemented with 20 µg ml^-1^ daptomycin (+ 50 µg ml^-1^ CaCl2) and survival of **(A)** SADR-1, SADR-2, SADR-1*^rpoB^*^_1^, and SADR-1*^clpP^*^_1^ or **(B)** JE2 and JE2 devoid of ClpXP activity due to expression of the ClpXI265E variant (described in the text) was followed by spotting 10 µl serially diluted cultures at the indicated time-points (*n* = 3) **(C)** Residual daptomycin activity in spent culture supernatants derived from indicated strains exposed to daptomycin for 6 h or 24 h was tested against SADR-1 cells (5 × 10^5^ CFU ml^-1^) spread on the surface of TSA plates. In both cases *n* = 3 in duplicates.

To understand how *S. aureus* cells could benefit from inactivation of ClpXP, we next determined the activity of daptomycin in spent supernatant derived from cultures of SADR-1, SADR-2, and the *rpoB* and *clpP* single mutants at 6 h and 24 h after addition of daptomycin to the bacterial cultures. The residual daptomycin activity was determined by using a zone of inhibition assay (47). After 6 h, daptomycin in spent supernatant from all strains was still active as observed by the efficient killing of *S. aureus* cells in the zone inhibition assay (Fig. 3C). Hence, inactivation of daptomycin activity does not explain the better survival of *S. aureus clpXP* cells at this time-point. We note, however, that daptomycin lost activity in spent supernatant from 24 h cultures of SADR-2, SADR-1*^clpP^*^_1^, and JE2 expressing the ClpX_I265E_ variant, while daptomycin from spent growth medium of the parental strains or SADR-1*^rpoB_^*^1^ remained active (Fig. 3C).

### The SNP in *clpP* counteracts the membrane-damaging defects associated with daptomycin exposure and makes cells adopt a rod-shaped morphology after daptomycin treatment

To understand how inactivation of ClpP promotes survival during daptomycin exposure, we next used super-resolution structured illumination microscopy (SR-SIM) to study the cell envelope in the four strains following exposure to 20 µg ml^-1^ daptomycin for 30 minutes or 3 hours using the same conditions as in the survival assay in Fig. 3. Exposure of *S. aureus* to daptomycin is associated with severe damage to the membrane and to follow integrity of the cell membrane, cells were stained with the nucleic acid stain, propidium iodide (PI, red) that is used as an indicator of severe membrane disruption because it is too large to penetrate intact membranes. In these experiments, the cell walls were also stained with Van-Fl, a fluorescent vancomycin derivative that stains the cell wall green by binding to the terminal D-Ala-D-Ala in non-cross-linked PG and with HADA (see below). As expected, PI-stained cells were not observed in cultures grown in the absence of daptomycin (Fig. 4). After 30 min exposure to daptomycin, the frequency of PI-positive cells was still very low in all strains. In contrast, the vast majority (> 90%) of SADR-1 and SADR-1*^rpoB^*^_1^ cells stained red after daptomycin treatment for 3 h, indicating severe membrane perturbation or even membrane collapse as suggested by the shrinkage in the size of many red-stained cells (Fig. 4A). At this time-point, PI-positive were also observed in cultures of SADR-2 and SADR-1*^clpP^*^_1^, however, the fraction of red cells was clearly reduced as compared to SADR-1 (Fig. 4A). Interestingly, many cells resisting PI staining appeared enlarged and had adopted a rod-shaped morphology and while all cells were spherical after 30 min daptomycin exposure, approximately half of SADR-2 and SADR-1*^clpP^*^_1^ cells were transformed into rods after 3 hours daptomycin treatment (Fig. 4A and 4B).

**Fig. 4.**
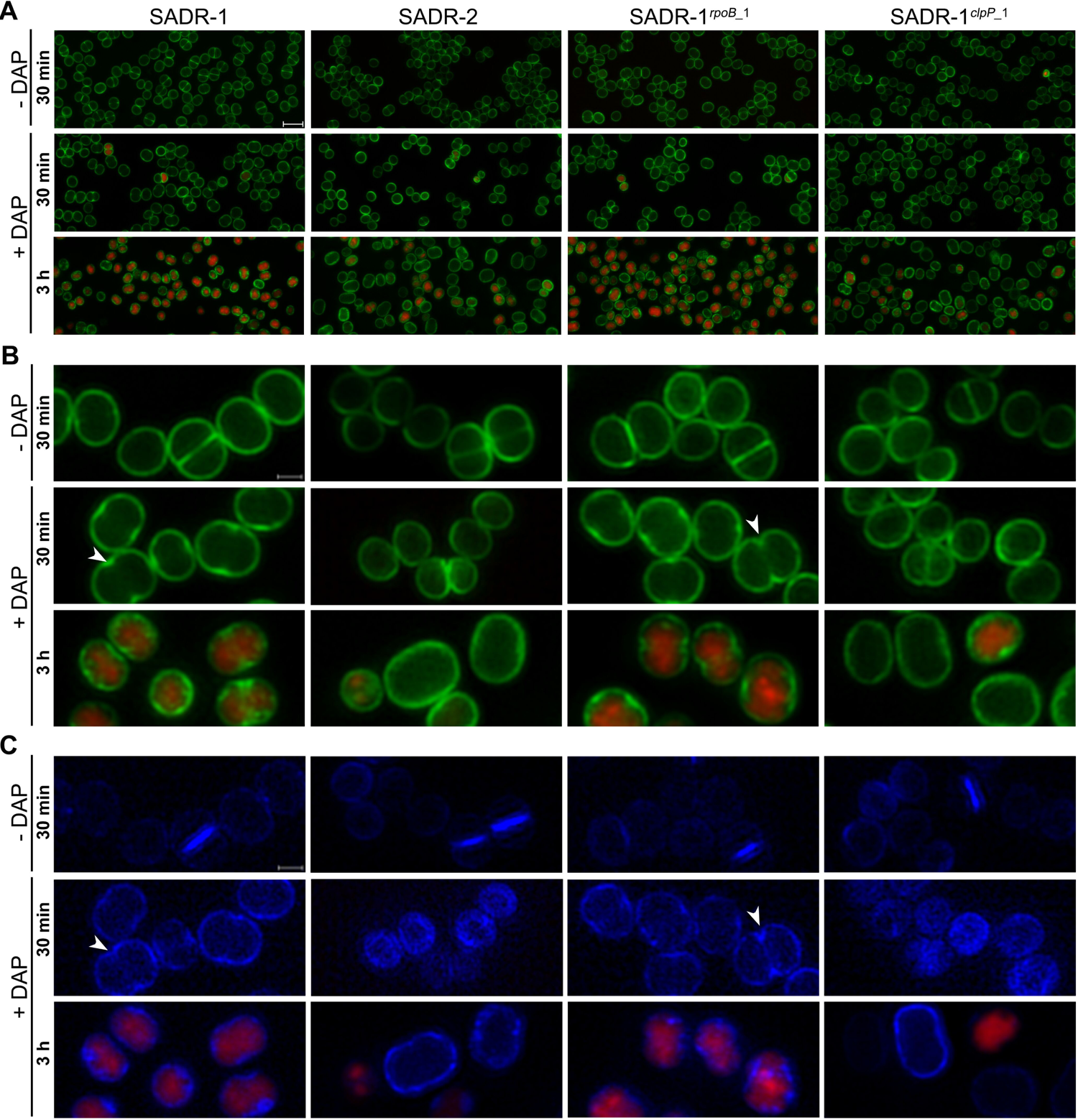
Inactivation of ClpP reduces PI-staining and promotes morphological changes following daptomycin treatment. Overnight cultures of the indicated strains were resuspended in TSB (∼2 × 10^8^ CFU ml^-1^) and incubated at 37°C ± 20 daptomycin for 30 min or 3 h before imaging with SR-SIM. Prior to SR-SIM, cells were labeled with PI (cells with compromised membranes, red), Van-FL (cell wall, green), and HADA (active cell wall synthesis, blue). **(A)** Upper panel shows overviews of cells stained with PI and Van-FL; scale bars, 2 μm. Middle and lower panels show enlarged cells (scale bar, 0.5 μm) of the four strains incubated ± daptomycin and stained with **(B)** Van-FL + PI, or **(C)** HADA + PI. The arrowheads point to cells with invaginations at mid-cell indicating premature splitting. The experiment was performed in duplicate with similar results.

To visualize membranes directly, SADR-1 and SADR-2 cells grown in the absence or presence of daptomycin were also stained with the membrane dye Nile Red prior to imaging. In the absence of daptomycin, the membrane stained uniformly red in both strains, however, following daptomycin exposure, membranes of SADR-1 cells stained irregularly with Nile Red, with the red signal being intensified in foci or patches along the cytoplasmic membrane (Fig. 5). In contrast, the membranes still had a uniform appearance in daptomycin treated SADR-2 cells (Fig. 5). This was also observed in SADR-1*^clpP^*^_1^ cells while membranes in SADR-1*^rpoB^*^_1^ cells appeared damaged as in SADR-1 cells (supplemental Fig. 1). We conclude that inactivation of ClpP abrogates the membrane damage induced by daptomycin and that *clpP* cells exposed to daptomycin tend to adopt a rod-shaped morphology.

**Figure 5.**
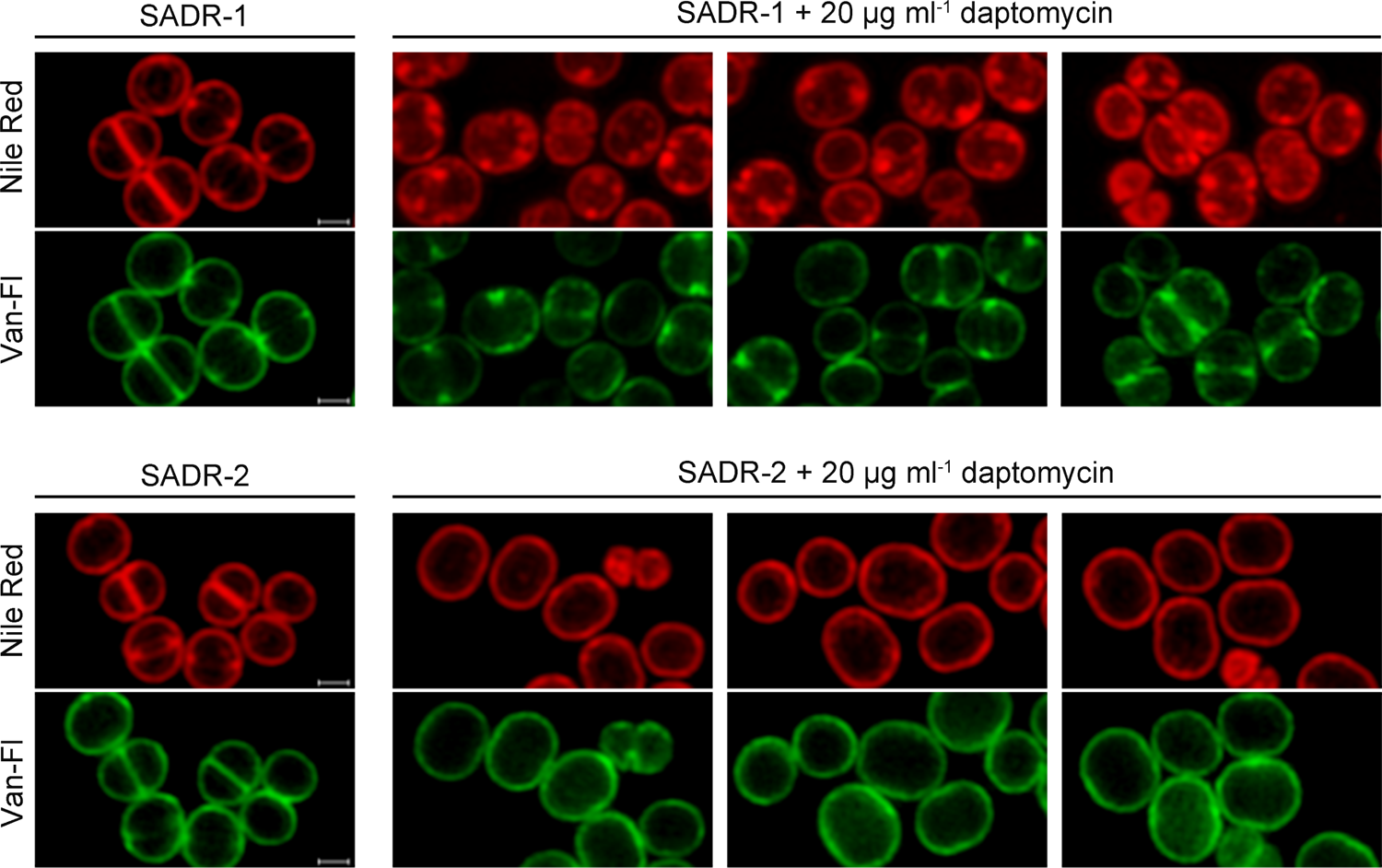
Reduced membrane damage in SADR-2 following daptomycin exposure. Overnight cultures of SADR-1 or SADR-2 were resuspended in fresh TSB (∼2 × 10^8^ CFU ml^-1^) and incubated at 37°C for 45 min in the absence of daptomycin, followed by incubation in TSB ± 20 µg ml^-1^ daptomycin at 37°C for 1 h. Prior to SR-SIM imaging, cells were labeled with Van-Fl (green) and membrane dye Nile Red (red). Scale bars, 0.5 μm.

### Shift of cell wall synthesis from the septal site to the peripheral wall in daptomycin-treated cells

The coccoid morphology of *S. aureus* depends on the balanced activity of two separate peptidoglycan (PG) synthesizing machineries operating in the outer wall and at the septal site, respectively (49). Depletion of the machinery essential for inward septum synthesis was shown by others to result in rod-shaped *S. aureus* cells (49), therefore, the appearance of rod-shaped SADR-2 cells following 3 h daptomycin exposure suggested that daptomycin alters the balance between these two machineries. To test this, regions of new PG insertion were visualized by incubating cells with the fluorescent D-amino acid, HADA (blue) prior to imaging (50). Indeed, this assay revealed that while untreated cells mainly incorporate HADA at the septal site, cells exposed to daptomycin for 30 min mainly incorporated HADA in the outer cell wall (Fig. 5C and supplemental Fig. 2). At this time point, the majority of SADR-1 and SADR-1*^rpoB^*^_1^ (> 85%) displayed a stronger HADA and Van-FL signal at mid-cell suggesting that cells have become synchronized at an early stage of septum formation (Fig. 5 and supplemental Fig. 2). We noted that some cells had an ‘hourglass’ shape (arrows in Fig. 5B and 5C), indicating that the premature septal ingrowths have been cleaved by autolytic enzymes (51, 52). This phenomenon was not observed in SADR-2 and SADR-1*^clpP^*^_1^ cells that displayed uniform Van-Fl staining of the outer cell wall after 30 min exposure to daptomycin (Fig. 5B). Strikingly, SADR-2 and SADR-1*^clpP^*^_1^ cells displayed uniform HADA-labeling of the outer cell wall even after 3 hours daptomycin exposure (Fig. 5C and supplemental Fig. 2). In conclusion, daptomycin seems to synchronize SADR-1 cells in an early stage of septum initiation with PG synthesis occurring both at the septal site and in the peripheral wall, while cells harboring the mutant variant of *clpP* continue PG synthesis only in the peripheral wall which could explain why cells transform from cocci to rods during the 3 h daptomycin exposure.

### Daptomycin binding is reduced in cells with inactivated ClpP

To visualize binding of daptomycin to the cell envelope of the four strains, cells were exposed to a mixture of Bodipy-FL-labelled daptomycin and unlabeled daptomycin for 30 min prior to imaging. Strikingly, SADR-2 and SADR-1*^clpP^*^_1^ cells were weakly stained with Dap-Fl as compared to SADR-1 and SADR-1*^rpoB^*^_1^ cells, indicating that inactivation of *clpP* reduces binding of daptomycin to the cell envelope of *S. aureus* (Fig. 6A). As described by others, the Dap-Fl signal was strongest at the septal site and in > 50% of SADR-1 and SADR-1*^rpoB^*^_1^ cells more intense Dap-Fl signals were observed in two foci at mid-cell corresponding to Dap-Fl being localized in a thin ring at mid-cell (supplemental Figure 3A). Notably, no invagination of the membrane was observed in such cells indicating that cells are in a very early stage of septation (Fig. 6B). Quantification of the Dap-Fl signal confirmed that the Dap-Fl signal was highly significantly reduced in SADR-2 as compared to SADR-1 (supplemental Figure 3). Monitoring of DAP binding over time revealed that prolonged incubation with daptomycin resulted in dispersion of Dap-Fl from the septal site in SADR-1 resulting in a uniform daptomycin signal throughout the entire cytoplasmic membrane at T=60 and bright foci throughout the cell at T= 90 (supplemental Figure 3). These observations confirm observations from a recent study reporting that daptomycin binding is biphasic with daptomycin binding occurring primarily at the septal site during the first phase followed by progressive dispersal of daptomycin in the entire membrane during the second phase eventually resulting in membrane collapse and cell shrinkage (17). In agreement with the improved survival of the SADR-2 cells, the weak Dap-Fl signal cells still localized uniformly to the cytoplasmic membrane in SADR-2 exposed to daptomycin for 90 min.

**Fig. 6.**
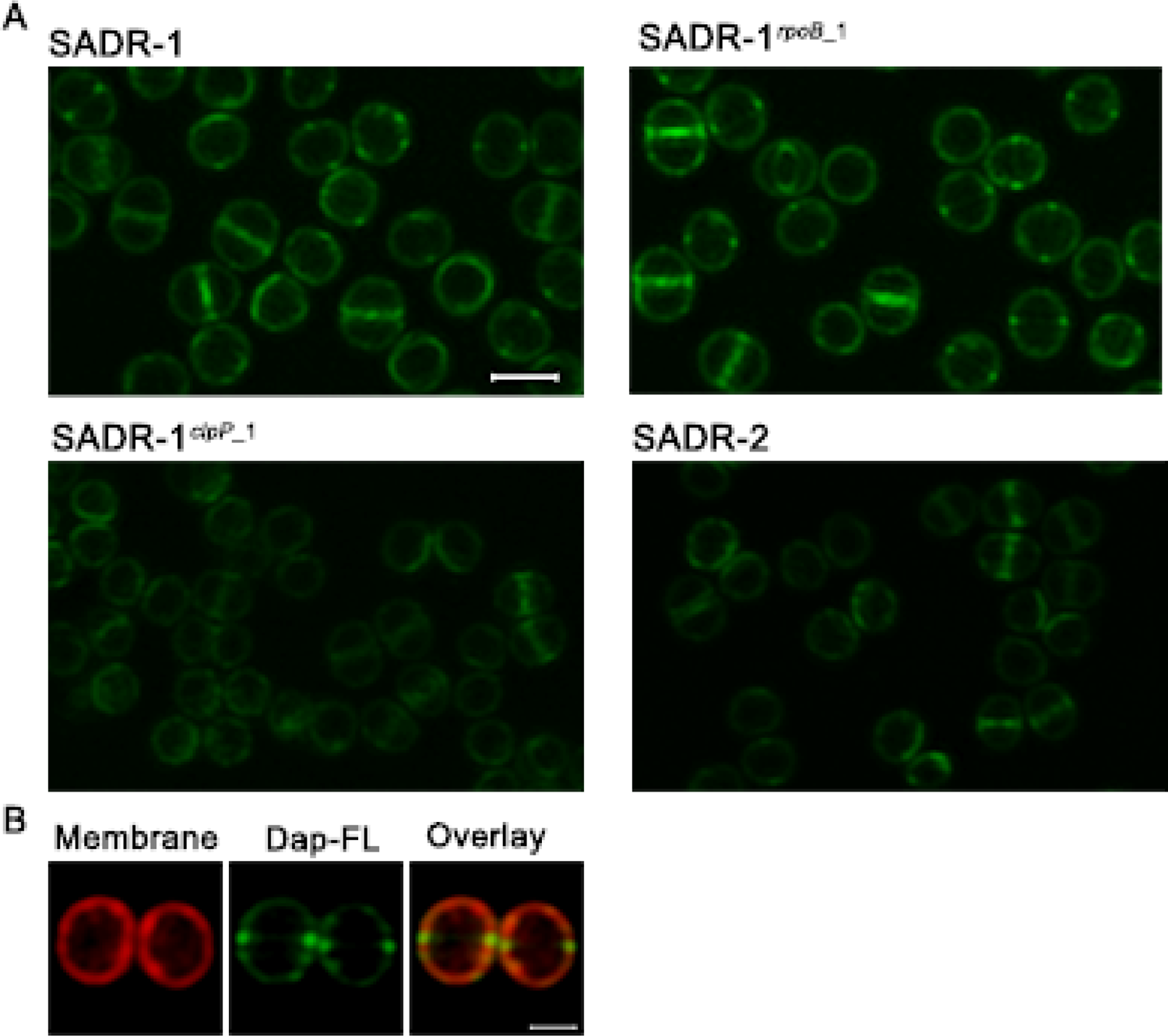
Inactivation of ClpP reduces binding of daptomycin to the cell envelope. (A) The four strains were grown to stationary phase (24 h) and incubated with a mixture of Dap-Fl and unlabeled daptomycin + Ca^2+^ for 30 min at 37°C before imaging by SR-SIM. Scale bar, 1 μm. (B) Nile Red staining of membranes showing that the membrane is not invaginated in cells with Dap-Fl signal in two foci at mid-cell. Scale bar, 0.5 μm.

### Daptomycin inhibits inward progression of septal PG synthesis in all strains

To more directly analyze how daptomycin interferes with spatiotemporal regulation of PG synthesis, we next sequentially labeled PG synthesis with two fluorescent D-amino acids of different colors: first, cells were labeled with HADA (blue) for 15 minutes in the absence of daptomycin. After washing away unbound HADA, cells were subsequently resuspended in TSB ± daptomycin and progression of PG synthesis was visualized by labelling for additionally 15 minutes with TADA (red) to create a virtual time-lapse image of PG synthesis in the presence or absence of daptomycin (Fig.7). Cells were imaged by SR-SIM, and progression of septal PG synthesis was analyzed by randomly picking SADR-1 and SADR-2 cells that had initiated septum synthesis during HADA-labeling and scoring cells according to the localization of the TADA signal (Fig. 7A). In the absence of daptomycin, septal TADA signal localized inwards to the HADA signal in > 90% of SADR-1 and SADR-2 cells, demonstrating that septal synthesis progresses inwards from the peripheral wall (Fig. 7). In daptomycin treated cells, however, the HADA and TADA signals instead co-localized in an early septal ingrowth in > 95% of SADR-1 and SADR-2 cells, a phenotype that was only observed in < 10% of non-treated cells (Fig. 7A). This finding supports that daptomycin interferes with inward progression of septal synthesis in both SADR-1 and SADR-2. Characterization of the *S. aureus* cell cycle has established that newly separated *S. aureus* daughter cells incorporate PG around the entire outer cell wall before initiating the next round of cell division (51, 53–54).

**Fig. 7.**
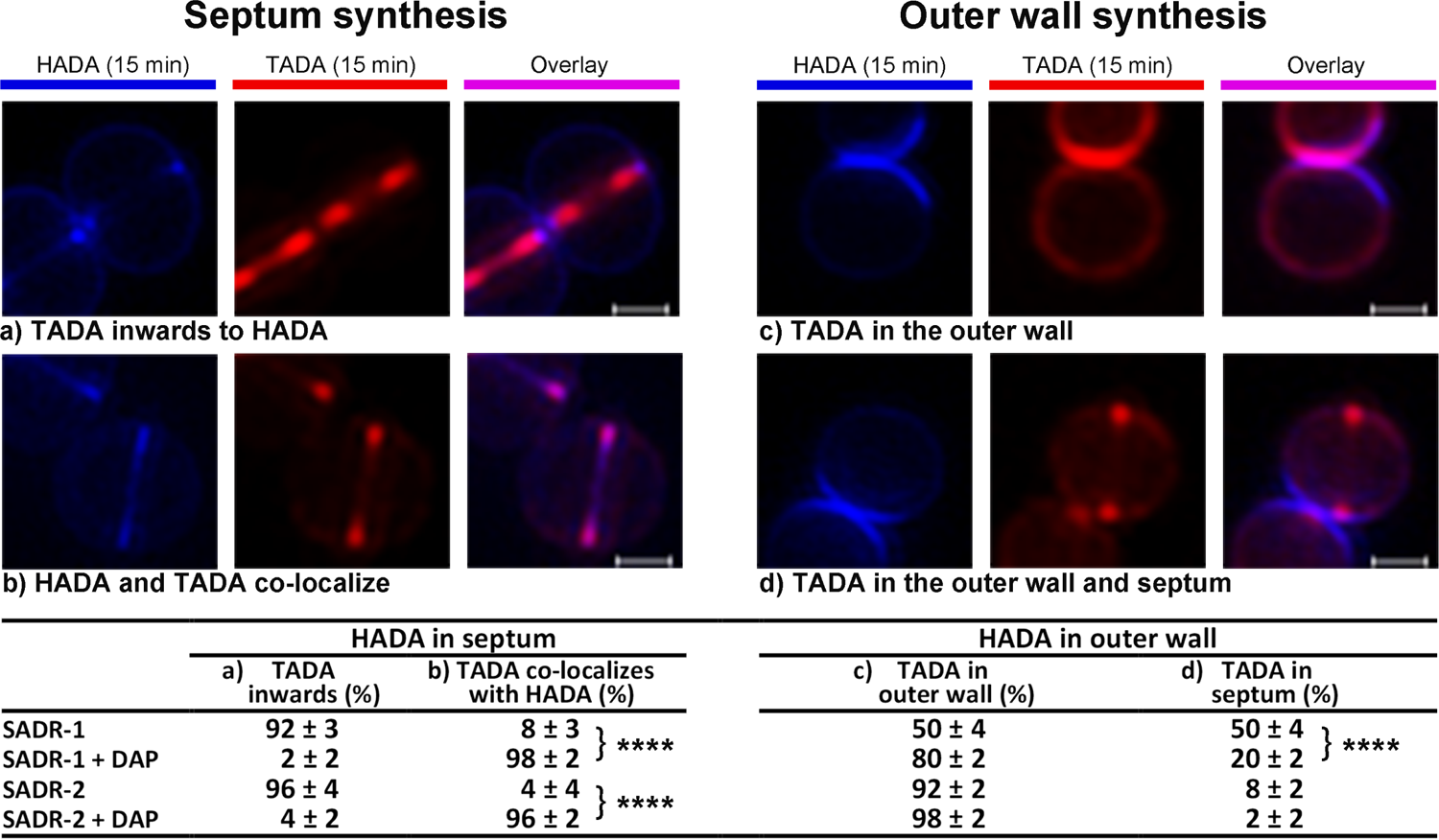
Daptomycin inhibits inward progression of septum synthesis and delays septum initiation. SADR-1 and SADR-2 cells (24 h cultures) were resuspended in TSB and incubated for 1 h at 37°C before cells were incubated with HADA (blue) for 15 min, followed by a washing step to remove unbound HADA. Subsequently, cells were resuspended in TSB (± daptomycin) and incubated with TADA (red) for 15 min. To access how daptomycin impacts PG synthesis at the septal site and in the outer cell wall, 50 cells for each condition displaying HADA signal, respectively, in early septal ingrowths, or, in the peripheral wall were randomly selected. Progression of PG synthesis was followed by assessing TADA incorporation as depicted in (**A-D**) and described in the text. Percentages are given as the mean and SD of three biological replicates. **** p < 0.0001; statistical analysis was performed using the Chi-square test for independence. Scale bars, 0.5 μm.

Consistent with this notion, the TADA signal localized in the entire peripheral wall of newly separated SADR-1 and SADR-2 cells that had completed septum synthesis during HADA-labeling (Fig. 7B). In the absence of daptomycin, approximately half of the newly separated SADR-1 daughter cells additionally displayed a TADA signal in early septal ingrowths, while TADA-stained septal ingrowths were observed in only 8% of newly separated SADR-2 cells, indicating that septum-initiation is delayed in SADR-2 as compared to SADR-1. In the presence of daptomycin, the TADA signal was still observed in the entire outer wall of newborn cells, however, the fraction of newborn cells displaying TADA-labeled septal ingrowth was reduced by more than two-fold in both strains (Fig 7B). Altogether, these results support the idea that daptomycin delays septum initiation and prevents inward progression of septum synthesis in both SADR-1 and SADR-2. In contrast, daptomycin does not seem to prevent PG-synthesis in the outer wall in either of the strains.

### Exposure to daptomycin upregulates cell wall hydrolases involved in daughter cell separation, and inactivation of ClpP mitigates this upregulation

In Fig. 5, we noted that some daptomycin-treated SADR-1 cells seemed to have initiated autolytic splitting of cells with premature septal ingrowths. Autolytic separation of *S. aureus* daughter cells relies on activity of the Sle1 amidase and the activity of the bi-functional Atl cell wall hydrolase (43, 55, 56), and we finally determined the amount of these two autolytic enzymes in cell wall extract derived before and after daptomycin exposure (45 min). Western blot analysis showed that the amount of Sle1 in the cell wall fraction increased approximately two-fold following incubation with daptomycin (Fig. 8). This increase was observed in all strains, however, consistent with Sle1 being a ClpP substrate (42, 43), the Sle1 levels are slightly elevated in the two strains, SADR-2 and SADR-1*^clpP^*^_1^, with inactive ClpP (Fig. 8A). Daptomycin exposure also increased Atl activity as shown by zymography (Fig. 8B). In zymograms, Atl-activity is observed in multiple bands reflecting that Atl is produced as a 138-kDa precursor protein containing a pro-peptide that is sequentially cleaved to generate 115- and 85-kDa intermediate products, and further processed to generate a 62-kDa *N*-acetylmuramyl-l-alanine amidase (AmiA), and a 51-kDa exo-β-N-acetylglucosaminidase (GlcA) that is not visible in zymograms (57). Following daptomycin exposure, the intensity of the 62-kDa AmiA band increased three-fold in cell walls derived from SADR-1 or SADR-1*^rpoB^*^_1^ cells, while the 62-kDa AmiA band did not increase in intensity in SADR strains harboring the mutant *clpP* allele (Fig. 8B).

**Fig. 8.**
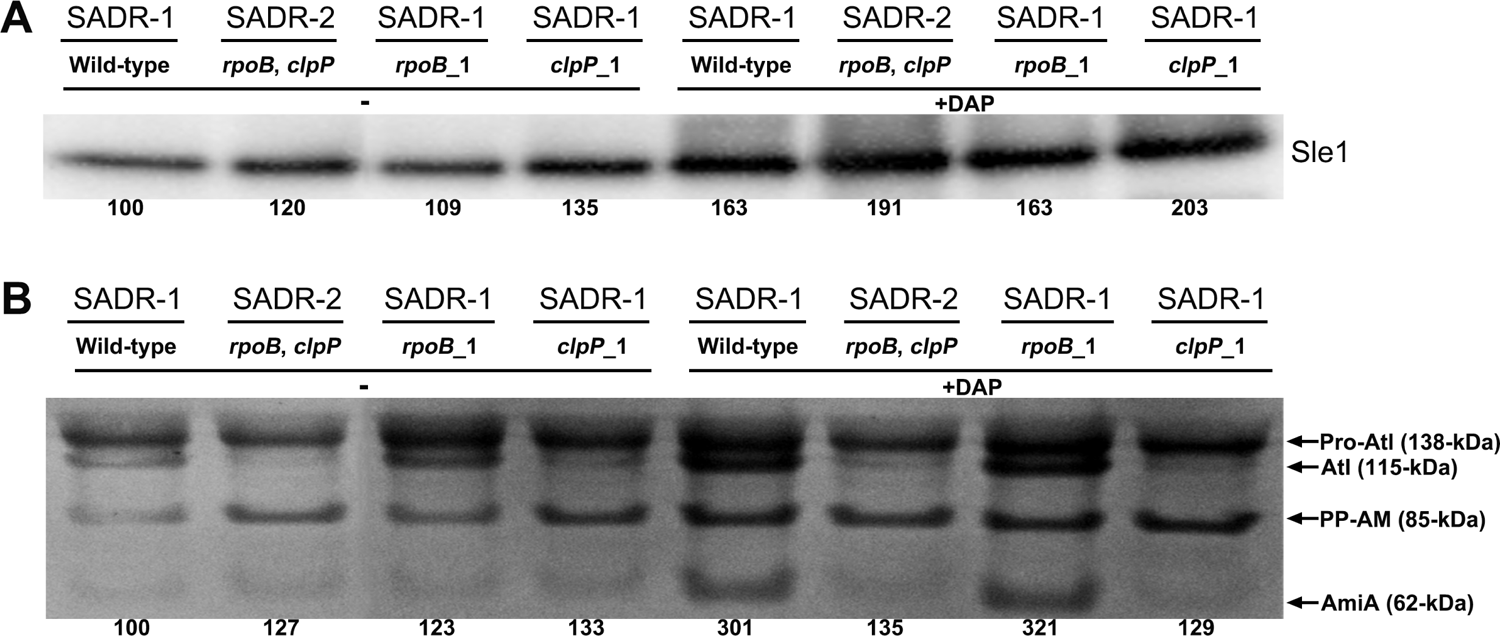
Daptomycin upregulates cell wall hydrolases involved in daughter cell separation. (A) Western blotting was used to determine the level of Sle1 in cell wall extracts derived from the indicated strains grown in the absence (-) or presence (+) of 0.4 µg ml^-1^ daptomycin + 50 µg ml^-1^ CaCl2 for 45 min. The Sle1 level was quantified using Fiji and normalized to the levels observed for the SADR-1 control. (B) Zymographic analysis performed with cell wall-associated proteins extracted from the indicated strains grown in the absence (-) or presence (+) of 0.4 µg ml^-1^ daptomycin + 50 µg ml^-1^ CaCl2 for 45 min. The sizes of the corresponding Atl bands were indicated on the right (in kilodaltons). The AmiA (62-kDa) activity was quantified by densitometric analysis using Fiji and further normalized to the levels observed for the SADR-1 control (grown in the absence of daptomycin).

Along this line, we noted altered processing of pro-Atl in *clpP* cells, with the 115-kDa Atl fragment (Atl without the pro-peptide) being diminished in *clpP* cells, both in the presence and absence of antibiotic. We conclude that daptomycin induces expression of the two major cell wall hydrolases involved in *S. aureus* daughter cell separation, and that inactivation of ClpP alters Atl processing and prevents induction of cell wall-associated AmiA following daptomycin exposure.

## Discussion

Here we show that a SNP in *clpP* acquired by an MRSA strain during daptomycin therapy *in vivo* eliminates ClpP activity resulting in well-described *clpP* phenotypes such as heat sensitivity, diminished expression of virulence factors, reduced cell size, and augmented tolerance to β-lactam antibiotics. We further show that the SNP in *clpP* increased the vancomycin and daptomycin MIC to the level of the clinical isolate collected after daptomycin treatment failure (MIC of 2 μg ml^-1^). This magnitude rise in MIC of daptomycin has been associated with therapeutic failure and poor patient outcomes (58), confirming the significance of the increase. Additionally, the mutation in *clpP* increased the abundance of more highly resistant subpopulations in daptomycin- and vancomycin PAP analyses. This is to the best of our knowledge the first demonstration that ClpP activity modulates resistance to the last-resort antibiotics vancomycin and daptomycin. The perhaps most important finding is, however, that the SNP in *clpP* also greatly promotes survival of *S. aureus* in the presence of high, therapeutic concentrations of daptomycin.

More than 30 years after the discovery of daptomycin, the killing mechanism of this first lipopeptide antibiotic in clinical use is still not fully elucidated (11). According to a recently proposed model (17), the Ca^2+^-DAP complex in the initial phase binds to the septal site where it forms a tripartite complex with the anionic phospholipid and cell wall precursors leading to a blockage of septum synthesis and delocalization of the septal cell wall biosynthetic apparatus. Consistent with this model, we observed immediate inhibition of inward progression of septal synthesis in daptomycin-exposed SADR-1 and SADR-2 cells (Fig. 7). As predicted by the model (17), this initial phase was followed by dispersal of daptomycin to the entire membrane, membrane disintegration and cell shrinkage in SADR-1. However, inactivation of *clpP* mitigated membrane damage which might be due to the reduced binding of daptomycin to the membrane in these cells, or, alternatively, the diminished membrane damage inhibits dispersal of daptomycin. Strikingly, daptomycin does not seem to inhibit PG synthesis in the outer cell wall, as cells continued to incorporate HADA in the outer cell wall both after short (30 min) and longtime exposure (3 h) to the antibiotic (Fig. 4; supplemental Fig. 2). While daptomycin exposed *clpP* cells continued PG synthesis uniformly in the peripheral wall eventually resulting in a rod-shaped morphology, daptomycin seemed to synchronize the parental strain in an early stage of septum synthesis with cell wall synthesis occurring in the lateral wall but being intensified at mid-cell (Fig. 4). We previously showed that arresting cells in an early stage of cell division may lead to premature autolytic splitting of the septal ingrowths (51, 52). Here we similarly observed splitting of the premature septal ingrowths in SADR-1, and we speculate that activation of septal autolysins in cells arrested in an early stage of septum synthesis may contribute to daptomycin imposed killing as depicted in the model in Fig. 9. At least two autolysins, Sle1 and Atl, contribute to *S. aureus* daughter cell splitting (43, 55,56). In support of our model, we observed that Atl activity was substantially increased in cell walls from daptomycin-exposed SADR-1 cells, while daptomycin exposure did not increase Atl activity in cell walls derived from cells with inactivated ClpP.

**Fig. 9.**
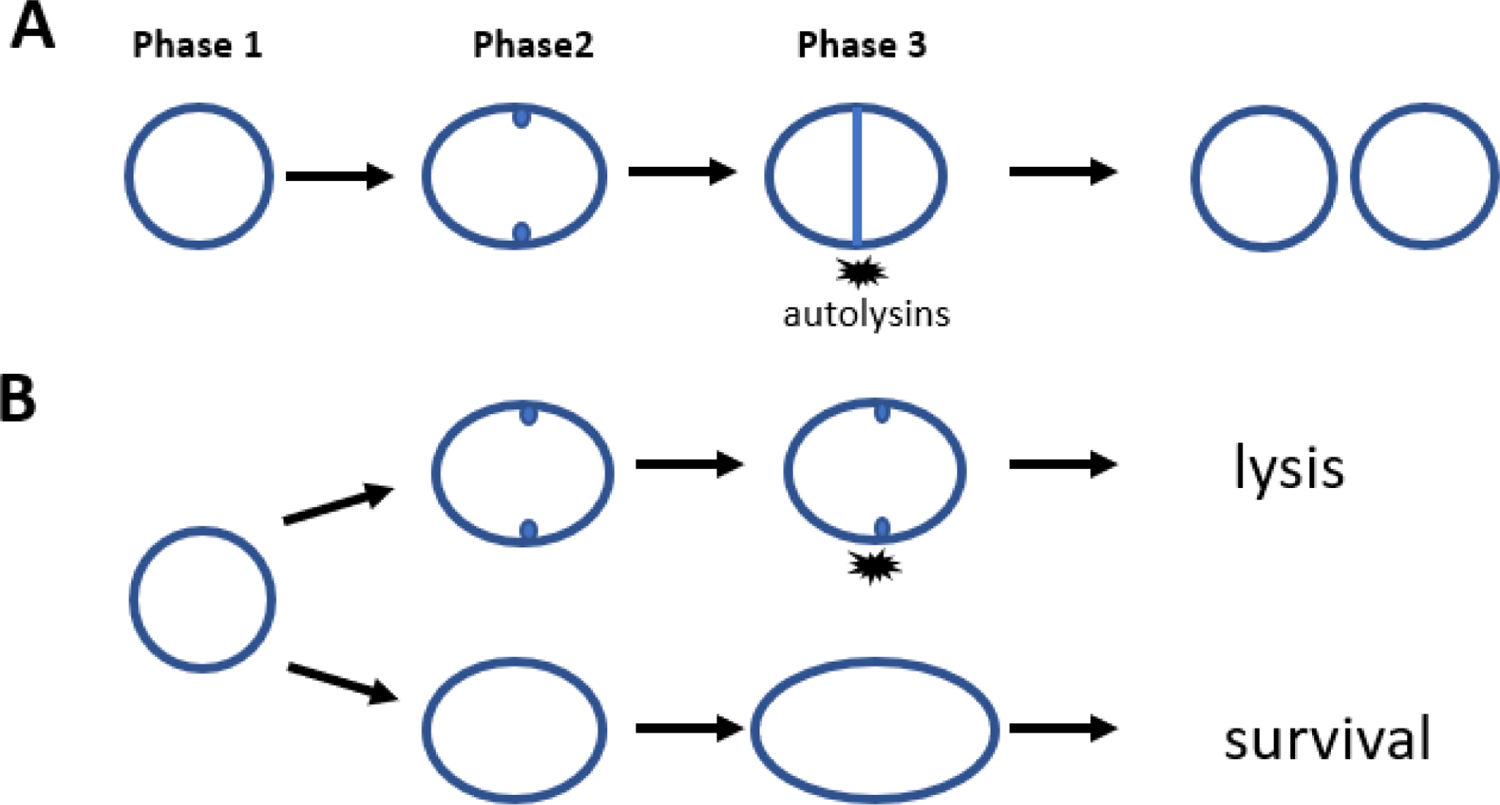
Proposed model. **(A)** The normal *S. aureus* cell cycle as described in (53): cells in phase 1 elongate prior to initiating septum synthesis; cells in phase 2 have initiated cell division, and cells in phase 3 have a closed septum. The cell cycle is completed when daughter cells separate by ultrafast popping initiated by cell wall hydrolases (autolysins). (**B**) In the presence of daptomycin, cells in phase 2 cannot complete septum synthesis and cells die from lysis elicited by the activation of septal autolysins. In contrast, cells in phase 1 survive because the septal autolysins are not activated. Instead, phase 1 cells continue elongating resulting in cells transforming into a rod shaped morphology that can resume cell division upon inactivation of daptomycin.

In contrast, surviving *clpP* cells seem to be arrested in the cell cycle prior to septum initiation which may protect cells from activation of septal autolysins. Daptomycin activity was abolished in spent supernatant from SADR-2 cultures incubated with daptomycin for 24 h (Fig. 3). At this time point, SADR-2 cells had resumed normal coccoid morphology (data not shown), indicating that rod-shaped cells are capable of dividing when daptomycin is no longer present.

The inactivating SNP in *clpP* was selected despite that it conferred heat sensitivity and diminished expression of central virulence factors such as Protein A, hemolysins, and extracellular proteases. This finding is in accordance with recent studies suggesting that *S. aureus* strains with low expression of toxins and other virulence factors are better adapted for long-term persistence and with studies showing that isolates with low-level resistance to daptomycin and vancomycin represent a bacterial evolutionary state favoring persistence over virulence (20, 59–61). The complex selective pressure *in vivo* that, in addition to the antibiotic treatments, involves the tissue microenvironments, and the immune responses, therefore, seems to favor mutations that at the same time render *S. aureus* less susceptible to the antibiotic and better equipped for survival in the host environment. Notably, using the same isolates as described here, we previously established that *clpP* inactivation may function as a potential immune evasion mechanism because ClpP is required for *S. aureus* to induce cell-surface expression of immune stimulatory NKG2D ligands on human monocytes (37). *S. aureus* infections occur in a variety of different tissue microenvironments that due to differences in nutrients, antibiotic exposures, immune responses, and spatial structure that will likely favor selection of site-specific adaptations. As an example, chronic osteomyelitis was recently associated with the ability of *S. aureus* to evade the immune system by invading the narrow osteocyte lacunocanalicular network (62, 63). Intriguingly, invading *S. aureus* transformed from 1 μm cocci to elongated rod-shaped cells of reduced diameter (62, 63). Hence, the small size of *S. aureus clpP mutant* and the daptomycin-imposed cell elongation could offer a selective advantage in some types of infections and therefore contribute to selection of such mutations *in vivo*. *In vitro* selection of mutants with reduced susceptibility to daptomycin has, however, also resulted in selection of a *S. aureus* strain with a substitution of a highly conserved glycine residue (G13) in ClpP, underscoring that *clpP* mutations, despite the associated fitness cost, can be selected by daptomycin alone (64).

Following daptomycin treatment failure, SADR-2 was isolated from the blood of a patient along with three other SADR-1 derivatives (8). The four strains all harbor the same SNP in *rpoB* strongly indicating that in response to daptomycin treatment SADR-1 first acquired the SNP in *rpoB* and then diversified. Strikingly, the SNP in *rpoB* was succeeded by three independent mutations that all relate to ClpXP-mediated proteolysis: as shown here, SADR-2 has acquired a loss-of-function mutation in the *clpP* gene, while two isolates (SADR-3 and SADR-4) harbor a frameshift mutation in *clpX* that abolishes ClpX expression (37), and, finally, a fourth isolate (A9798) has a frameshift mutation in *yjbH* encoding a presumed adaptor protein selecting certain substrates for degradation by ClpXP (8, 65). So far, the essential transcriptional regulator Spx is the only known substrate recognized by YjbH (65–67) and accordingly we previously found Spx to accumulate in SADR-2, 3, and 4 (36). The independent selection of mutations that inactivate all subunits of the ClpXP-YjbH protease complex is indicative of a strong selective pressure for stabilizing Spx. In *B. subtilis* the Spx thiol-stress regulon is also induced in response to heat stress and cell wall stress (68, 69) and we speculate that constitutive expression of the Spx regulon may contribute to survival of *S. aureus* during antibiotic-imposed stress in the cell envelope. Experimental validation of this hypothesis is, however, challenged by the essentiality of the Spx protein in *S. aureus* (67).

In conclusion, *S. aureus* may benefit from inactivating ClpXP during *in vivo* therapy because bacterial cells become better equipped for surviving antibiotic treatments and to evade host responses. The pathways underlying better survival are likely complex and multi-faceted.

## Materials and methods

### Strains and culture conditions

The *S. aureus* strains used in this study are listed in Table 1. Unless otherwise stated, SADR-1 and its derivatives were cultivated at 37°C in 20 ml tryptic soy broth (TSB; Oxoid) in 200 ml Erlenmeyer flasks with linear shaking at 180 rpm or on tryptic soy agar (TSA; Oxoid) plates. The growth of *S. aureus* strains was assessed by measuring optical densities at a wavelength of 600 nm (OD_600_). In all experiments, bacterial strains were freshly streaked from the frozen stocks on TSA and incubated overnight at 37°C. From these plates, TSB cultures were inoculated to an OD_600_ of <0.05.

### Antibiotic susceptibility testing

Antibiotic susceptibility testing (MICs of daptomycin, vancomycin, oxacillin and rifampin) was carried out by the Danish National Reference Laboratories for Resistance Surveillance (SSI, Copenhagen, Denmark) using Etest strips (bioMérieux) or The Sensititre™ Vizion™ broth microdilution system (Thermo Fisher Scientific) using the strain *S. aureus* strain ATCC29213 as a reference strain. All antibiotic susceptibility testing of strains was performed in biological triplicate.

### Population analysis profiles (PAPs)

Antibiotic resistance profiles were determined as previously described (36). Overnight cultures of tested strains were normalized to an OD_600_ of 1.0 and serially diluted to 10^-6^ in 0.9% NaCl. 100 µl of appropriate dilutions were spread on TSA plates supplemented with increasing concentrations of vancomycin, oxacillin, or daptomycin + 50 µg ml^-1^ CaCl_2_. The numbers of CFU ml^-1^ were determined after 48 h of growth at 37°C to allow for growth of slow-growing subpopulations.

### Bacterial survival in the presence of high concentrations of daptomycin

Survival in the presence of high concentrations of daptomycin was performed as previously described (47): ∼2 × 10^8^ CFU from stationary cultures (18 h) were resuspended in 6 ml TSB supplemented with 20 µg ml^-1^ daptomycin + 50 µg ml^-1^ CaCl_2_ and incubated at 37°C with linear shaking (180 rpm). Bacterial survival was determined by spotting appropriate dilutions (10^1^ to 10^5^-fold) of cultures on TSA plates at the indicated time points. The plates were incubated at 37°C for 24 h before counting CFU.

### Daptomycin activity in spent supernatant

The residual daptomycin activity in spent supernatant was determined as described previously (47): at the indicated times, 1 ml of culture was removed and supernatants were collected by centrifugation (17000 g, 10 min) followed by sterile-filtration (0.22 µm filter). Sterility of the supernatant was confirmed by spotting 30 µl of the supernatant on a TSA place and incubating for 24 h at 37°C. Wells of 10 mm were made in TSA plates containing 50 µg ml^−1^ CaCl_2_ followed by the spreading of 100 µl stationary SADR-1 cells (∼10^6^ CFU ml^−1^ in TSB) across the surface. The spread bacterial inoculum was allowed to air dry before the wells were filled with 200 µl spent culture supernatant. Plates were incubated for 18 h at 37°C before evaluating daptomycin activity.

### Western blot analysis

To determine the level of Sle1 and Protein A in *S. aureus* cells, the bacterial strains were grown in TSB at 37°C with aeration until the OD_600_ reached 1.0. At this point, 1 ml of cells from each strain was harvested, and an extract of total cellular proteins was prepared by harvesting the cells by centrifugation and resuspending cell pellets in 50 mM Tris-HCl (pH 8.0) (200 µl per OD unit) and incubating with 5 µg ml^-1^ lysostaphin (Sigma) for 30 min at 37°C. To determine the amount of Sle1 in cell walls of *S. aureus* grown in the absence or presence of daptomycin, exponentially growing cultures (OD_600_ = 0.8) were divided into two cultures that continued growth in the absence or presence of 0.4 µg ml^−1^ daptomycin + 50 µg ml^−1^ CaCl_2_ for an additional 45 min. Cell wall associated proteins were purified from 15 ml of culture after washing cells once with 10 ml cold 0.9% NaCl. To release proteins from the cell wall, cells were incubated with 4% SDS (1 ml per OD unit) for 45 min at 25°C with gentle shaking. Cells were precipitated by centrifugation and the supernatant containing the cell wall associated proteins was collected. 20 µl of each sample was loaded on NuPAGE 4-12% Bis-Tris gels (Invitrogen), and electrophoresis was performed according to the manufacturer’s instructions. After separation, proteins were blotted onto a PVDF membrane (Invitrogen) using an XCell II Blot Module system (Invitrogen). To detect Sle1, the PVDF membrane was firstly blocked with human IgG to evade a signal from protein A. The Sle1 protein was probed using rabbit anti-staphylococcal Sle1 antibody at a 1:2500 dilution. To detect protein A, the membrane was probed using an anti-Sle1 antibody without pre-blocking with human IgG and cellular proteins extracted from a *spa* mutant were used to indicate the position of the protein A signal. Detection of the specific protein signal was achieved by using WesternBreeze Chemiluminescent (anti-rabbit) kit (Invitrogen).

### Zymographic analysis

Bacteriolytic enzyme profiles were obtained as described (70) using a 10% SDS-PAGE containing 0.1% (w/v) heat-inactivated SA564 (methicillin-sensitive CC5 isolate) as substrate. Cell wall-associated proteins were prepared as described for western blotting. Following electrophoresis, the gel was washed with deionized water for 45 min and then incubated in renaturation buffer (50 mM Tris-HCl (pH 7.5), 1% Triton X-100, 10 mM CaCl_2_ and 10 mM MgCl_2_) at 37°C for 20 h. The gel was stained with staining buffer (0.4% methylene blue 0.01% KOH, 22% EtOH) for 1 min, and de-stained with deionized water for 5 h.

### SR-SIM analysis

i. *Image acquisition*. For SR-SIM analysis, images were acquired using SR-SIM with an Elyra PS.1 microscope (Zeiss) using a Plan-Apochromat 63x/1.4 oil DIC M27 objective and a Pco.edge 5.5 camera. Images were acquired with five grid rotations and reconstructed using ZEN software (black edition, 2012, version 8.1.0.484) based on a structured illumination algorithm, using synthetic, channel-specific optical transfer functions and noise filter settings ranging from −6 to −8. Laser and staining specifications can be found in Table 3. Prior to imaging cells were placed on an agarose pad (1.2% in PBS). All SR-SIM analysis was performed at the Core Facility of Integrated Microscopy (CFIM).
ii. *Estimation of cell volume*. Prior to imaging, cultures of *S. aureus* were grown at 37°C with shaking (180 rpm) until OD_600_ ∼0.4. Cells were stained for 5 min with the membrane dye Nile Red and were visualized by SR-SIM, as described above. The volume of 100 phase 1 cells (cells without ingrowing septa) was determined as previously described (53). Briefly, the cell shape of *S. aureus* was assumed to be that of a prolate spheroid, and the volume was estimated using the equation V = 4/3πab^2^, where *a* and *b* correspond to the major and minor axes, respectively. An ellipse was fitted to the border limits of the membrane to acquire measurements of the major axis (a) and minor axis (b). Ellipse fitting and measurements were carried out using Fiji software (https://imagej.net/Fiji). The statistical analyses were performed using RStudio software (version 4.2.2). A chi-squared test of independence was used to determine whether there was a significant relationship between daptomycin exposure and PG progression at the septal site or in the outer cell wall under the tested condition. A P value of < 0.05 was considered significant.
iii. *Determining the morphological changes induced by daptomycin*. To determine morphological changes induced by daptomycin, ∼2 × 10^8^ CFU from stationary cultures (18 h), were collected and washed once in 0.9% NaCl. Cells were collected and suspended in 6 ml fresh TSB in the absence or presence of 20 μg ml^-1^ daptomycin + 50 μg ml^-1^ CaCl_2_. Cells were incubated at 37°C for 30 min or 3 h before imaging with SR-SIM. Prior to imaging, cells were stained for 5 minutes at 37°C using the following dyes: PI (cells with a compromised membrane), Van-fl (cell wall), and HADA (active PG synthesis).
iv. *Determination of PG progression of S. aureus*. To determine the PG progression, ∼2 × 10^8^ CFU from stationary cultures (18 h) of SADR-1 or SADR-2 were collected and washed once in 0.9% NaCl before suspending cells in 6 ml fresh TSB and growing for 1 h at 37°C. The cells were then sequentially labeled with FDAAs of different colors; cells were initially incubated with HADA (blue) for 15 min, washed twice in PBS and resuspended in TSB supplemented with TADA (red) in the absence or presence of 20 μg ml^-1^ daptomycin + 50 μg ml^-1^ CaCl_2_. After labeling for 15 min, the cells were washed twice in PBS and imaged as described above. The progression of PG synthesis at the septal site and in the outer cell wall was accessed by randomly picking 50 cells of each condition displaying HADA signal, respectively, in early septal ingrowths, or, in the peripheral wall. Progression of PG synthesis was followed by assessing the location of TADA signal. Percentages are given as the mean and SD of three biological replicates.
v. Visualizing membrane damage induce by daptomycin. To visualize the membrane damage induced by daptomycin, ∼2 × 10^8^ CFU from stationary cultures (18 h) of the indicated strains were collected and washed once in 0.9% NaCl before suspending cells in 6 ml fresh TSB and growing for 45 minutes at 37°C in the absence of daptomycin followed by incubation at 37°C for 1 h in the absence or presence of 20 µg ml^-1^ daptomycin + 50 μg ml^-1^ CaCl_2_. Prior to imaging, cells were stained for 5 min at room temperature with Van-Fl (cell wall) and Nile Red (membrane). Images were acquired using the Elyra PS.1 as previously described.
vi. *Determination of daptomycin binding*. To study daptomycin binding in SADR-1 and its derivatives, cells were grown for 18 h at 37°C in TSB with aeration. At this time, ∼2 × 10^8^ CFU were collected and washed once in 0.9% NaCl before suspending cells in 6 ml fresh TSB supplemented with a mixture of labeled and unlabeled daptomycin to a final concentration of 20 μg ml^−1^ (18 μg ml^−1^ unlabeled Dap and 2 μg ml^−1^ Dap-FL) and 50 μg ml^-1^ CaCl_2_. (Dap-FL was kindly provided by Professor Tanja Schneider, University of Bonn). Cells were incubated for 5, 30, 60, or 90 minutes. To visualize the cell membrane, Nile Red was added when indicated and samples were incubated for 5 minutes at room temperature. After staining, cells were washed twice in 0.9% NaCl. To quantify the amount of bound Dap-FL, the integrated density (intDen) was measured. Integrated density is equivalent to the product of Area and Mean Gray Value, meaning that it takes the difference in cell size into account. The integrated density was determined in at least 800 random cells for each sample. Analysis was performed on cells only stained with Dap-FL to avoid any leakage from other channels in samples with low Dap-FL signal and consequently, it was not possible to distinguish between different stages in the cell cycle or to objectively exclude possible dead cells which gives a much stronger signal. Measurements were carried out in Fiji and analyzed using GraphPad Prism 9.5 (GraphPad Software LLC). An unpaired, two tailed t-test was used to compared differences in daptomycin binding.

## Acknowledgments

We greatly acknowledge Professor Tanja Schneider (Bonn University) for providing Dap-Fl and Motoyuki Sugai (Hiroshima University) for donating the Sle1 antibodies. A very special thanks to the staff at the Core Facility for Integrated Microscopy (University of Copenhagen) for their enthusiastic assistance in doing microscopy.

---

**Supplemental Fig. 1.** Inactivation of *clpP* mitigates daptomycin-imposed membrane damage. The indicated strains were grown to stationary phase (24 h) and exposed to 20 µg ml^−1^ daptomycin + Ca^2+^ for 60 min at 37°C before staining of membranes with Nile Red and imaging by SR-SIM. Representative pictures are shown from three independent experiments. Scale bar, 1 µm.

**Supplemental Fig. 2.** Daptomycin shifts cell wall synthesis from the septal site to the peripheral cell wall. Overnight cultures of the indicated strains were resuspended in TSB (∼2 × 10^8^ CFU ml-1) and incubated at 37°C ± 20 µg ml^-1^ daptomycin for 30 min or 3 h before imaging with SR-SIM. Prior to SR-SIM, cells were labeled PI (cells with compromised membranes, red), Van-FL (cell wall, green), and HADA (active cell wall synthesis, blue). Overview pictures showing HADA-labeling of the cells shown in Fig. 4 before and 30 min and 3 h following daptomycin exposure.

**Supplemental Fig. 3.** Quantification of Dap-Fl binding in SADR-1 and SADR-2 at different time points. Approximately 2 × 10^8^ CFU ml^-1^ of the indicated strains (18 h cultures) were incubated with a mixture of labeled and unlabeled daptomycin to a final concentration of 20 μg ml^−1^ at 37°C before imaging with SR-SIM. Prior to SR-SIM at the indicated time points. Overview pictures showing Dap-Fl-labeling of the cells at the different time points are shown to the left. At T=30 and T= 60, the integrated density was determined in at least 800 random cells for each sample (right side). Measurements were carried out in Fiji and analyzed using GraphPad Prism 9.5 (GraphPad Software LLC). Scale bar, 1 µm.

**Supplemental Table 1.**
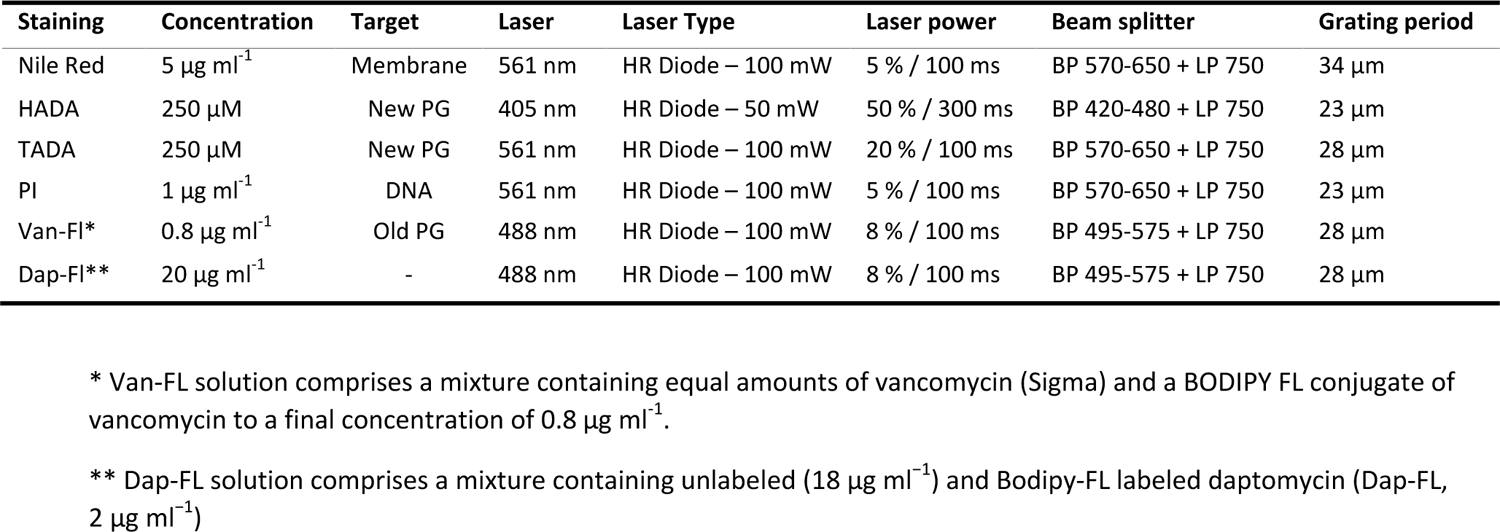
Staining and laser specifications used for SR-SIM

| Staining | Concentration | Target | Laser | Laser Type | Laser power | Beam splitter | Grating period |
| --- | --- | --- | --- | --- | --- | --- | --- |
| Nile Red | 5 $\mu\text{g ml}^{-1}$ | Membrane | 561 nm | HR Diode – 100 mW | 5 % / 100 ms | BP 570-650 + LP 750 | 34 $\mu\text{m}$ |
| HADA | 250 $\mu\text{M}$ | New PG | 405 nm | HR Diode – 50 mW | 50 % / 300 ms | BP 420-480 + LP 750 | 23 $\mu\text{m}$ |
| TADA | 250 $\mu\text{M}$ | New PG | 561 nm | HR Diode – 100 mW | 20 % / 100 ms | BP 570-650 + LP 750 | 28 $\mu\text{m}$ |
| PI | 1 $\mu\text{g ml}^{-1}$ | DNA | 561 nm | HR Diode – 100 mW | 5 % / 100 ms | BP 570-650 + LP 750 | 23 $\mu\text{m}$ |
| Van-FL* | 0.8 $\mu\text{g ml}^{-1}$ | Old PG | 488 nm | HR Diode – 100 mW | 8 % / 100 ms | BP 495-575 + LP 750 | 28 $\mu\text{m}$ |
| Dap-FL** | 20 $\mu\text{g ml}^{-1}$ | - | 488 nm | HR Diode – 100 mW | 8 % / 100 ms | BP 495-575 + LP 750 | 28 $\mu\text{m}$ |

\* Van-FL solution comprises a mixture containing equal amounts of vancomycin (Sigma) and a BODIPY FL conjugate of vancomycin to a final concentration of 0.8 $\mu\text{g ml}^{-1}$ .
\*\* Dap-FL solution comprises a mixture containing unlabeled (18 $\mu\text{g ml}^{-1}$ ) and Bodipy-FL labeled daptomycin (Dap-FL, 2 $\mu\text{g ml}^{-1}$ )

